# Characterization of a novel amber-reassigned Crassvirales genus infecting *Segatella copri* from Egypt

**DOI:** 10.64898/2026.08.21.746148

**Authors:** Lamyaa M. Ibrahim, Marwa T. ElRakaiby, Mohamed H. Habib, Hamdallah H. Zedan, Tamer A. Mansour

## Abstract

Bacteriophages of the order Crassvirales are currently believed to be the most prevalent dsDNA phages in the human gut virome, yet their global biogeography and genomic diversity remain poorly characterized due to an overrepresentation of industrialized Western studies in public repositories. In this study, we integrated computational metagenomics and molecular approaches to identify and validate the first complete Crassvirales genome from an Egyptian population. De novo assembly and viral profiling yielded a 101,034 bp circular genome (contig k141_108779) predicted to infect the non-industrialized gut symbiont *Segatella copri*. The genome displays the notable feature of *amber* stop codon reassignments (NCBI Genetic Code 15), where canonical (TAG) stop codons encode glutamine (Q). This alternative code increases coding density to 91%. Population-level PCR surveillance and Sanger dideoxynucleotide sequencing across 252 individual Egyptian fecal samples, pooled in 10 composites, confirmed the active circulation and local sequence heterogeneity of this lineage within the community. Phylogenomic and intergenomic similarity analysis demonstrated that the isolate shares less than 50% total average nucleotide identity with all recognized type strains. These data establish that this phage constitutes a novel species within a newly proposed genus inside the family *Darmviridae*. Our findings expand the known geographic distribution of crAss-like phages, highlight translational versatility among *Segatella*-infecting viruses, and emphasize the importance of expanding virome cohorts to underrepresented regions.

## 1. Introduction

The vast majority of viruses on Earth, often referred to as “viral dark matter,” remain uncharacterized due to limitations in laboratory culturing and the absence of a universal marker gene analogous to ribosomal DNA in bacteria ^1,2^. The human gut virome is a prime example of an ecosystem dominated by these unexplored viral entities, and viral metagenomics provides an unparalleled opportunity for their discovery ^3^. Among the most significant contributors to the stability of the gut ecosystem are bacteriophages, which coexist with their bacterial hosts in a delicate balance.

A remarkable breakthrough in viral metagenomics was the computational discovery of a widespread human gut bacteriophage, now known as prototypical crAssphage (p-crAssphage). It was identified through the re-analysis and cross-assembly of public fecal metagenomes, where it was found to be six times more prevalent than all other identified phages combined^3^. This initial discovery has since expanded to a vast and diverse group of related viruses, termed crAss-like phages, which infect bacteria of the phylum Bacteroidota, primarily the family Bacteroidaceae.

These phages are now classified under their own taxonomic three orders, Crassvirales, Metacrassvirales and Paracrassvirales, which comprise seven families, 75 genera and numerous species, established based on total average nucleotide identity, shared orthologous genes, core-proteome and single-gene phylogenetic analysis^4^. Despite their widespread prevalence, with some studies showing they comprise up to 90% of an individual’s gut viral reads^3,5,6^, the biogeography of Crassvirales remains underexplored in many parts of the world, including Egypt. It is unclear whether populations in these regions harbor distinct species of crAss-like phages compared to those found in industrialized communities.

This study aimed to bridge this knowledge gap by investigating the existence of crAss-like phages in an Egyptian gut microbiome study. Using a systematic approach that combined the re-analysis of public metagenomic data with computational genomics and molecular validation, we identified and characterized a novel Crassvirales genome and assessed its genomic relationship to the currently classified members of crAss-like phages.

## 2. Materials and Methods

### 2.1 Public Metagenomic Data and Assembly

Publicly available fecal shotgun metagenomic data from Egypt were explored using the HumanMetagenomeDB server (release 1.1; last accessed December 2025)^7^. The only available dataset, comprising fecal samples from 28 Egyptian teenagers (SRA accession ERR726368), was downloaded from the NCBI Sequence Read Archive^8^. Sequencing reads were already quality-filtered, trimmed, and stitched into single fragments. The stitched reads were assembled into contigs using Megahit v1.2.9 with default parameters for single-end reads^9^.

The quality of the assembly was evaluated for contiguity and completeness using MetaQUAST v5.3.0 and CheckM2 v1.0.2+galaxy1, respectively^10,11^. Contigs were binned for CheckM2 by MetaBAT2 v2.17+galaxy0 using default settings^12^. In addition, the assembly correctness was assessed by quantifying the proportion of reads incorporated into the assembly by mapping input reads back to the assembled contigs using Bowtie2 v2.5.4 in end-to-end mode, and the number of aligned reads was determined using SAMtools v1.22^13,14^.Another metric of correctness is the average percentage identity of assembled contigs aligned against a database of reference sequences for 145 crAss-like phages downloaded from the NCBI Nucleotide repository. These genomes are the sequences used by the International Committee on Taxonomy of Viruses (ICTV) to define the three orders of Metacrassvirales, Paracrassvirales, and Crassvirales (Taxonomy Proposal 2025.013B.Crassvirales_reorganisation).^4^ Also, they include all (n =74) the RefSeq reference sequences of Crassvirales (last checked in May 2026).^15^

### 2.2 Viral Sequence Prediction and Identification of crAss-like Phages

Viral sequences were predicted using three tools: VirSorter v2.2.4+galaxy0, VIBRANT v1.2.1+galaxy2, and ViralVerify v1.1^16–18^. Circularity of contigs was determined by detecting terminal redundancy (>130 bp at ≥97% identity) using BLASTn tool v2.14.1+^19^. Finally, the putative viral contigs of crAss-like phages were identified according to the criteria of the ICTV for the order *Crassvirales* published in 2021 (Taxonomy Proposal 2021.021B.A.Crassvirales)^20^ and augmented by features advised by others ^21,22^ as follows: Contigs were selected to exceed 10 kb in length and contain no ambiguous bases (relaxed from the 50kb cutoff suggested by ICTV to increase the sensitivity of our initial screen), exhibit viral signatures scores >0.5 with DeepVirFinder v1.0 and *p*-values <0.05 and are classified as “complete” or “high-quality” by CheckV v1.0.3^23,24^. Furthermore, candidate sequences were required to show >70% BLASTn identity over an alignment length ≥3 kb and an *E-value* <1e−10 to known crAss-like phages, while having coding density >80% using either the translation genetic table 11 or 15.

### 2.3 Genomic Characterization and Functional Annotation

The final candidate genome was analyzed to detect the viral genomic termini and packaging mode using PhageTerm v1.0.12^25^. Bacterial host prediction was performed using the CHERRY tool as part of PhaBOX v2.0 web-server with a CHERRYScore > 0.9, and lifestyle (lytic/temperate) was inferred using BACPHLIP (score >0.8)^26–28^.

Genome annotation was carried out in two steps. First, the annotation module of Cenote-Taker3 v3.4.1 was executed with the options -p F -am T to identify possible open reading frames and their possible functional annotations^29^. Second, amino acid sequences of hypothetical proteins were further annotated using HHpred^30,31^ against multiple databases: *PDB_mmCIF70_25_May*^32^, *Pfam-A* v37.0^33,34^, NCBI-CDD v3.19^35^, and *TIGRFAM* v15.0^36^ while applying default settings. Hits with a probability ≥80% and coverage >30% were considered for downstream functional annotation. GC skew, replichores, and circular genome map visualizations were made on the CGView (Proksee) Server^37^. The tRNA genes were predicted using tRNAscan-SE v2.0 with bacterial settings (-B)^38^.

### 2.4 Alternative Genetic Code Analysis

Prodigal v2.6.3 was used to compare annotations with the standard genetic code (11; TAG → stop), and alternative codes 4 (UGA → W) and 15 (TAG → Q)^39^. Coding density was defined as the total length of predicted genes divided by the genome length, and an alternative genetic code was inferred if coding density increased by >5% relative to predictions with the standard code. In addition, codon usage was reported as the percentage of a specific codon out of all codons in a coding region of consecutive windows. Calculations of codon usage were done in consecutive windows of 500bp using a python script^40,41^

### 2.5 Phylogenomics and Comparative Genomics

A phage proteomic tree based on genome-wide similarity was generated using ViPTree v4.0 to compare the newly assembled phage with the 145 crAss-like phages^42^. Average Amino Acid identity (AAI) was calculated using EzAAI v1.2.4^43^.Amino acid sequences of four core proteins (large terminase subunit, major capsid protein, portal, DNA primase) highly conserved across the three orders, in addition to the Egyptian phage, were aligned by MAFFT v7.526 (online service) using the L-INS-i algorithm^44^. For each protein, only the longest annotated sequence from each genome was retained. Then, multiple alignments for each protein were curated using the ClipKit v2.1.3 Kpic method to select informative sites^45^. Afterwards, they were manually curated before phylogeny reconstruction using the browser version of JalviewJS, where truncated sequences were removed; also, numerous inteins were removed from alignments by identifying and removing the columns present in <10% of individual sequences^46^. Maximum likelihood trees and optimal models were inferred using IQTree v3.1.1 (options --alrt 1000 -B 1000 -T AUTO)^47–49^. The core-proteome maximum likelihood phylogenetic tree of the three orders was based on concatenated protein alignments of the four core proteins. The concatenation of these alignments by AMAS^50^ was used to build the maximum likelihood tree with IQTree (options -allnni -nm 4000 -bb 1,000 -alrt 1,000). All trees were midpoint rooted, and the Cellulophaga phage phi13:2 (NC_021803) was used as an outgroup. For orthologous gene sharing, amino acid sequences of ORF protein products were clustered into orthologous groups using OrthoFinder v3.1.0^51^. Then, shared orthologous percentages were calculated for all 145 crass-like phages and the Egyptian phage using a python script^52^. For heatmaps, genomes were clustered hierarchically using the complete linkage method with Euclidean distances, based on the shared orthologous percentages matrix (implemented in ClustVis v1.0 web tool)^53^.

Furthermore, to assess the novelty of the assembled phage, BLASTn was used to query the assembled genome against the standard nucleotide BLAST database (nr/nt) (May 2026 release). Sequences showing >80% genome coverage and >80% identity were tested for intergenomic similarities (tANI), in addition to members of the *Darmviridae* family, against the Egyptian phage using the VIRIDIC web service^54^.

### 2.6 Molecular Validation in Egyptian Fecal Samples

No ethical approval was required for fecal sample collection. This study followed the ethical guidelines of the Faculty of Pharmacy, Cairo University, and was approved by its Scientific Research Ethics Committee (approval serial number: MI 3041, 2021).

Fecal samples for PCR analysis were collected from 252 Egyptian individuals participating in a national health campaign in Giza Governorate. Samples were pooled and homogenized into ten composites (Supplementary Table S13). DNA extraction was done using the QIAamp Fast DNA Stool Mini Kit, following the large sample (>220 mg) protocol.

PCR primers were designed using FastPCR to target specific regions of the assembled phage genome^55^. PCR amplification reactions (20 µL) contained 10 µL of 2× amaR onePCR mix, 0.5 µM each of forward and reverse primers, and 20 ng of template DNA. Thermal cycling conditions were: initial denaturation at 94 °C for 3 min; 30 cycles of 94 °C for 30 s, 60 °C for 1 min, and 72 °C for 2 min; followed by a final extension at 72 °C for 5 min. Amplicons were purified using the EasyPure PCR Purification Kit and sequenced by Macrogen (Seoul, South Korea) to confirm identity.

## 3. Results

### 3.1 Metagenomic Assembly and Identification of a Novel Crassvirales Genome

Only one gut shotgun metagenomic dataset was found in HumanMetagenomeDB server ^7^. This comparative study involved 28 healthy male teenagers (13.9 ± 0.6 years old (mean ± s.d.)) (Supplementary Table S1). Assembly of the Egyptian metagenome (8.8 million single-end reads, average length 242 bp) generated 389,741 contigs with a total assembly size of 305.4 Mb (305,407,776 bp) and an average GC content of 45.33%. It had an N50 contig length of 869 bp and a contig size ranging from 200 bp to 119684 bp with an average of 783 bp. Assembled contigs were grouped into 27 MAGs and according to CheckM2 and MIMAG standards, only one suited the criteria for high quality (≥90% completeness, <5% contamination), eight were medium quality (≥50% completeness, <10% contamination), and the remaining 18 were classified as low quality (<50% completeness, <10% contamination) (Supplementary Table S4)^11,56^. Backmapping of the input reads to the final assembled contigs resulted in a ∼74% remapping rate. In addition, alignment of assembled contigs to 145 crAss-like phages published by the ICTV (Taxonomy Proposal 2025.013B.Crassvirales_reorganisation)^4^(Supplementary Table S2) identified 1169 alignments, of which 148 were above 1000 bp and had an average identity of 88.63%, confirming the successful reconstruction of crAss-like viral sequences from the sequencing data (Supplementary Table S3).

Our screening pipeline (Figure 1) integrated three complementary viral prediction tools and applied stringent filtering criteria to identify high-quality crAss-like phage genomes with the least possible contamination by non-phage sequences. This computational workflow identified 1621 putative dsDNA viral sequences. Only 11 viral contigs longer than 10,000 bp were predicted by all three tools (Supplementary Table S5, S6, S7). Further filtration using DeepVirFinder retained eight sequences with scores >0.5 and p-values <0.05 (Supplementary Table S8)^23^. Remarkably, BLASTn alignments of these 8 contigs against the 145 crAss-like phage reference genomes identified one contig, referred to as k141_108779, with substantial similarity to 8 distinct members of the order Crassvirales fulfilling the BLASTn criteria of the ICTV report for putative viral contigs belonging to crAss-like phages (Taxonomy Proposal 2021.021B.A.Crassvirales) (Supplementary Table S3)^4,20,57,58^. Its best match belonged to the newly classified family *Darmviridae*, with the highest query coverage (44,765 bp) and similarity (88.29%) to Smehivirus intestinihominis, uncultured phage cr29_1 (E-value = 3.53E^-29^) (Supplementary Table S9). CheckV analysis identified a direct terminal repeat (DTR), confirming that contig k141_108779 represents a complete, circularized viral genome (Supplementary Table S10)^24^.

**Figure 1:**
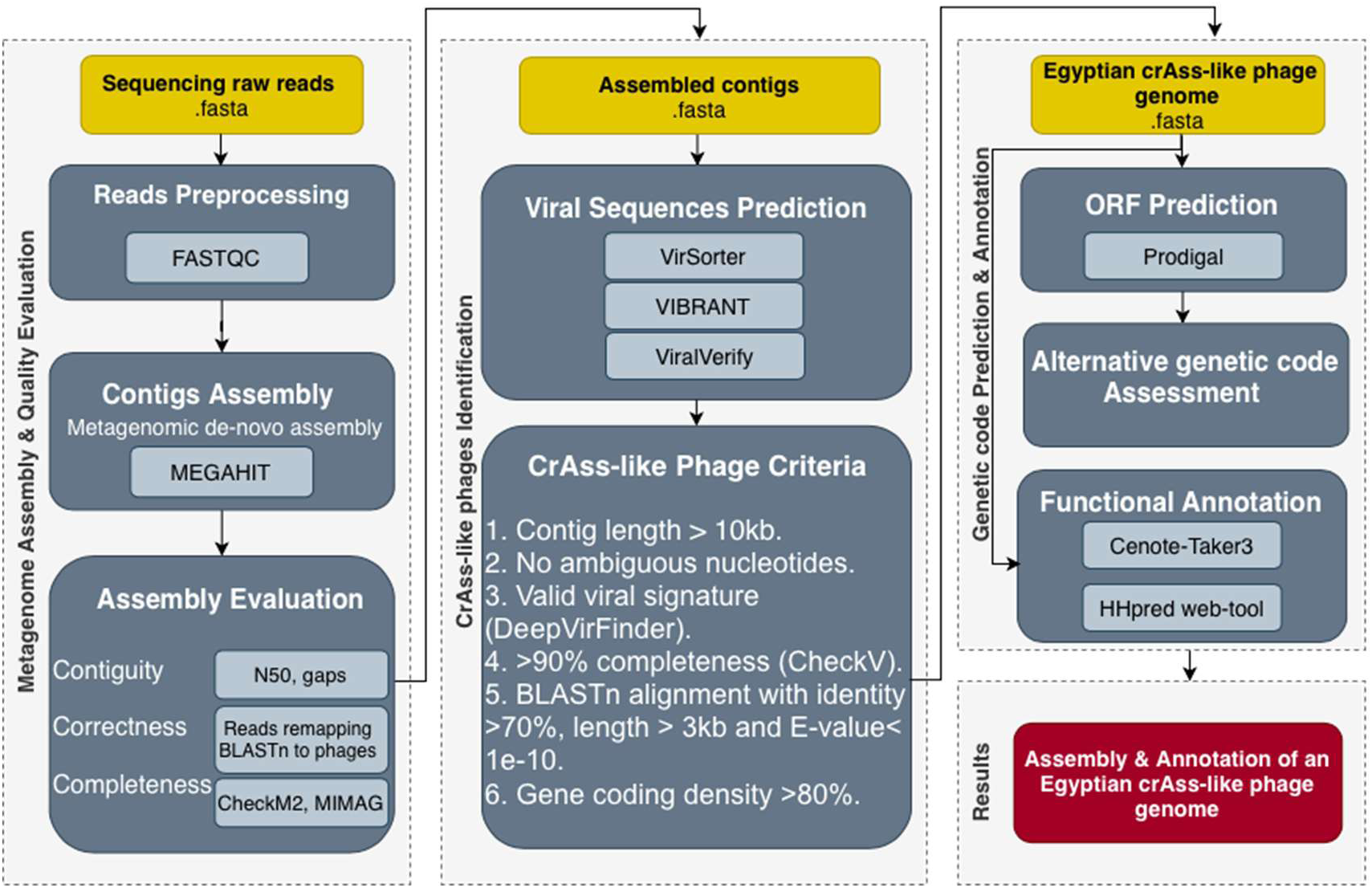
Bioinformatics workflow for identification and functional annotation of the Egyptian crAss-like phage. Yellow boxes: input data (publicly available trimmed and joined reads). Grey boxes: computational tools and processes (parameters described in Methods). The final output is a fully annotated phage genome.

### 3.2 Genomic Characterization and Host-Lifestyle Inferences

The Egyptian crAss-like phage possesses a circular genome of 101,034 bp with 36.43% GC content. Analysis of local read coverage revealed circularly permuted, redundant phage genome termini with headful DNA packaging and terminase initiation occurring at a *pac* site. CHERRY predicted *Segatella copri* (formerly *Prevotella copri*) as the primary host with a high CHERRYScore of 0.94, a result supported by a CRISPR-based matching method that identified sequence homology with the host’s adaptive immune record^28^. Simultaneously, BACPHLIP evaluated the phage’s proteome for 206 lysogeny-associated protein domains using a Random Forest classifier^26^. The analysis identified several conserved domains (cd01123, cd03408, pfam03050, pfam04855, and pfam18803), yet the specific pattern and absence of critical temperate markers—such as integrases and excisionase—resulted in a 0.9986 probability of a virulent lifestyle (Supplementary Table S14). These high-confidence predictions suggest the Egyptian phage functions as a lytic predator of *S. copri*, a key human gut symbiont.

### 3.3 Reassignment of the Amber (TAG) Stop Codon

A notable observation of this phage genome was its unusually low coding density (∼59%) when genes were predicted using the standard bacterial genetic code (code 11). Under this model, many predicted genes appeared abnormally short or fragmented, with internal stop codons, especially the canonical TAG *amber* stop codons, truncating what should have been a single, long protein (Grey ring, Figure 2)^59,60^. Notably, many of these short ORFs encoded fragments of conserved structural proteins, including the portal protein and the terminase large subunit (Supplementary Figure 1), as well as essential metabolic proteins like RNA polymerase, endonuclease, and dsDNA-binding domain. In contrast, frequency analysis of codon usage revealed a systematic avoidance of *amber* (TAG) stop codons across other large genomic regions (orange bars, Figure 2). Together, these observations strongly suggest the use of an alternative genetic code in which canonical stop codons are reassigned to encode amino acids rather than terminate translation.

**Figure 2:**
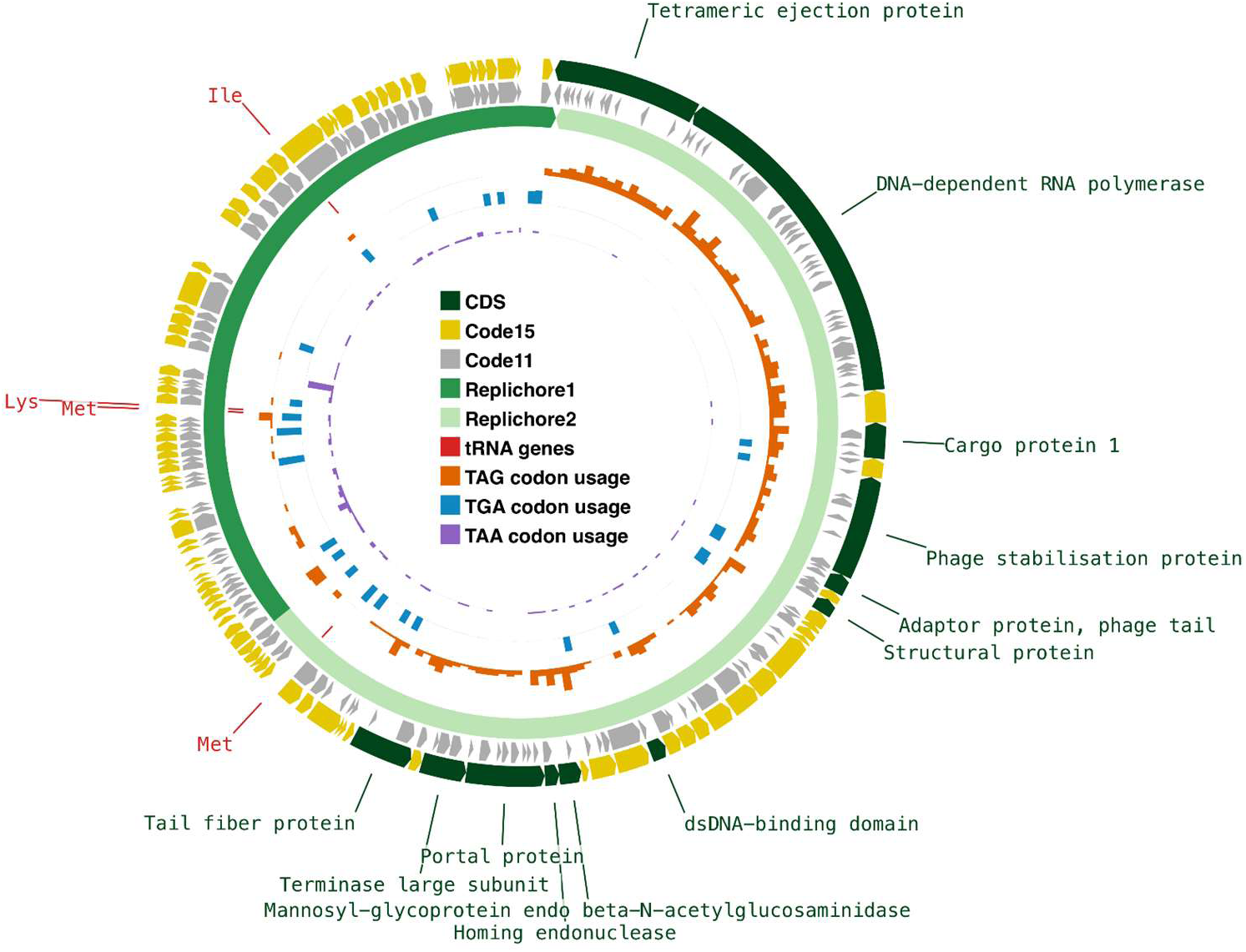
Manually curated genomic map of the Egyptian phage after reassigning TAG stop codons to glutamine (NCBI genetic code 15). Dark green: structural, lysis and metabolic genes enriched with in-frame TAG codons (orange bars). Yellow: DNA replication genes using standard code (TGA/TAA stops, blue and purple bars). Alternating green shades indicate replichores; GC skew suggests bidirectional replication.

Consistent with this hypothesis, re-annotation using NCBI genetic code 15 where the TAG stop codon is repurposed to encode glutamine (Q), would transform fragmented genes into single, full-length, functional proteins, including TerL, portal, and RNAP subunits^61^. This re-annotation substantially increases the overall coding density to ∼91%, aligning with values typically observed in members of Crassvirales and Paracrassvirales orders (Yellow ring, Figure 2, Supplementary Figure 2, Supplementary Table S11. These predicted genes are arranged in two opposite orientations separated by an inverted GC skew, indicative of two replichores (alternating shades of green, Figure 2).

### 3.4 Functional gene annotation

Functional gene annotation was performed using a two-step approach. Initially, the Cenote-Taker3 pipeline, designed for viral genome discovery and annotation, identified 30 genes^29^. Cenote-Taker3 uses pyhmmer to run hmmscan of a single protein query sequence against its curated database of viral Hidden Markov Model (HMMs)^62,63^. Also, the pipeline invokes mmseqs2 to identify viral hallmark genes and known functional domains by homology search against the Conserved Domain Database(CDD)^35,64^. Next, HHpred was utilized to examine the remaining hypothetical proteins (it is specifically designed to detect remote homology based on pairwise comparison of profile HMMs)^30,31^. This approach enabled the annotation of another 27 genes and the classification of an additional 12 genes into conserved protein families with unknown functions. Altogether, 102 genes were identified, including 27 involved in replication and metabolism, 7 transcription regulatory genes, 11 structural genes, 4 packaging and assembly genes, 6 tRNA genes, one lysis gene, and 13 genes were assigned to families of unknown functions. The remaining genes encode hypothetical proteins. (Figure 3, Supplementary Table S12). No genes associated with a lysogenic life cycle, such as an integrase or excisionase, were found, supporting the prediction of a virulent lifestyle.

**Figure 3:**
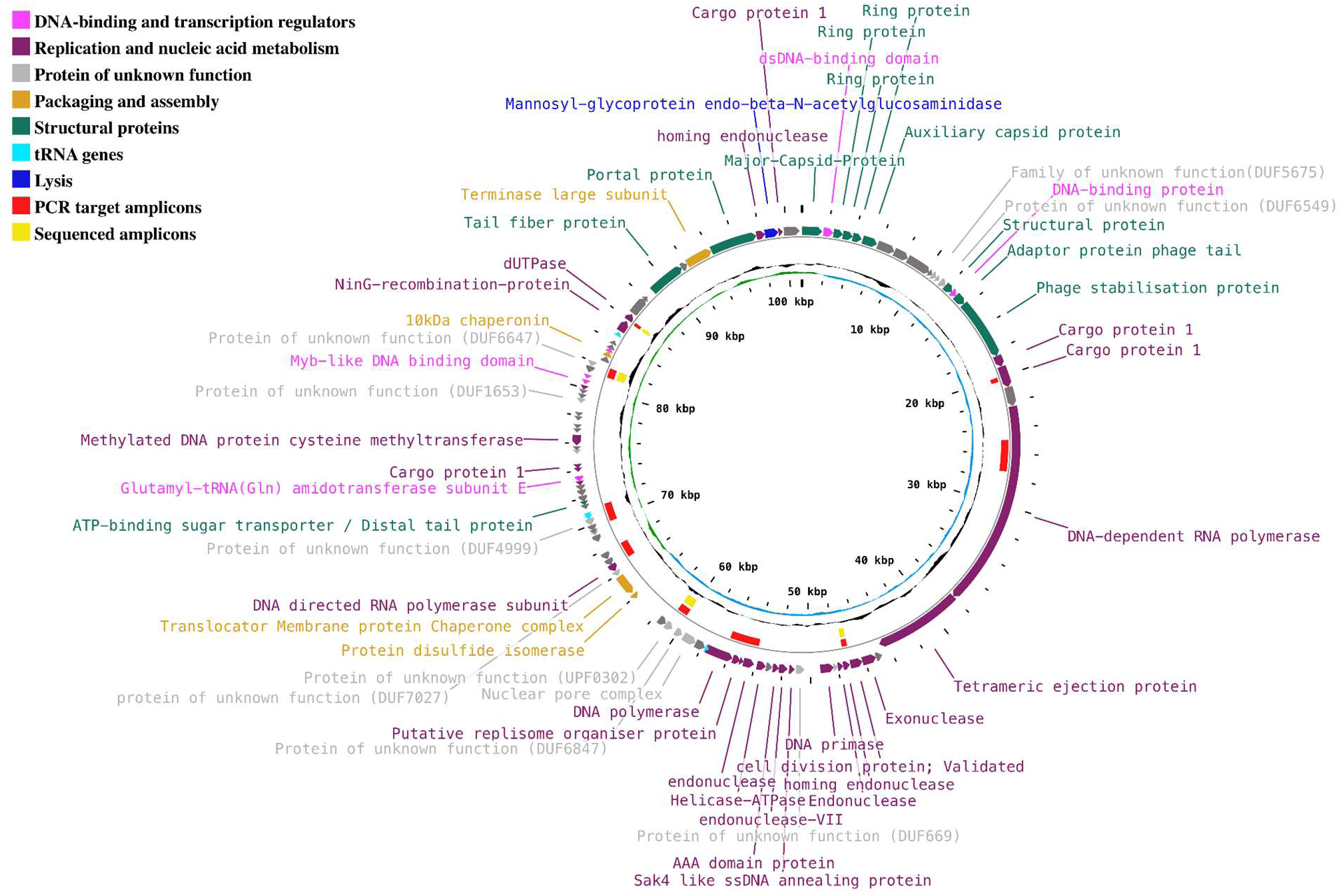
Circular genome map of the Egyptian crAss-like phage. Inner circle: GC skew (positive/negative). Middle circle: G+C content. Outer circles: protein-coding genes (ORFs) coloured by functional category (see legend). Yellow segments: PCR amplicon targets. Red bars: regions confirmed by Sanger sequencing

Although stop-codon-reassigned phages frequently encode suppressor tRNAs to facilitate translational readthrough^40,60,65^, no suppressor tRNA with a matching CUA anticodon was detected. Instead, the genome encodes six tRNA loci, comprising four high-confidence functional tRNAs (two specific for methionine, one for lysine, and one for isoleucine), with predicted lengths and isotype scores (49.8–74.6) consistent with functional tRNAs. The remaining two putative pseudogenes exhibited low isotype scores (25.9–26.1), indicating incomplete or ambiguous tRNA-like structures. Notably, annotation reveals the presence of a glutamyl-tRNA(Gln) amidotransferase, which likely supports mechanisms involving TAG stop codon reassignments into glutamine.

### 3.5 Molecular Screening Confirms Phage Presence in the Egyptian Population

To validate the computational assembly and confirm the circulation of this virus within the local population, we collected 252 fecal samples from several health units in Giza: Manial Shiha, Zawya Abou Muslim, Shubra Ment, and Tersa (Supplementary Figure 3). Samples covered a wide age range [106 early young (<12 years old), 43 young (13-20 years old), 46 early adults (21-40 years old), 33 middle-aged (41–60 years old), and 24 elderly (≥61 years old)]. This was done as part of the Egyptian national campaign for the early detection of schistosomiasis, fascioliasis, and intestinal parasites, which commenced between June and October in 2023 and again in 2024^66^. After collection, samples were pooled into ten composites (Supplementary Table S13).

Ten sets of PCR primers were designed to target the discovered phage genome, and four successfully amplified products of the expected size from DNA extracted from pooled fecal samples (Figure 4, Table 1). This provides molecular evidence that our phage circulates within the wider Egyptian community. Sanger sequencing of these amplicons confirmed their identity, showing 94-99% sequence similarity to the assembled reference genome (Supplementary Figure 4). Subsequent BLASTn alignments of these amplicon sequences against members of the three orders of crAss-like phages confirmed the novelty of amplicon U1 sequence with only 32% coverage and 75.7% similarity to uncultured crAss-like phages (Supplementary Figure 5). Meanwhile, amplicon U2 shows a distant relationship with 82% coverage and ∼82% nucleotide identity.

**Figure 4:**
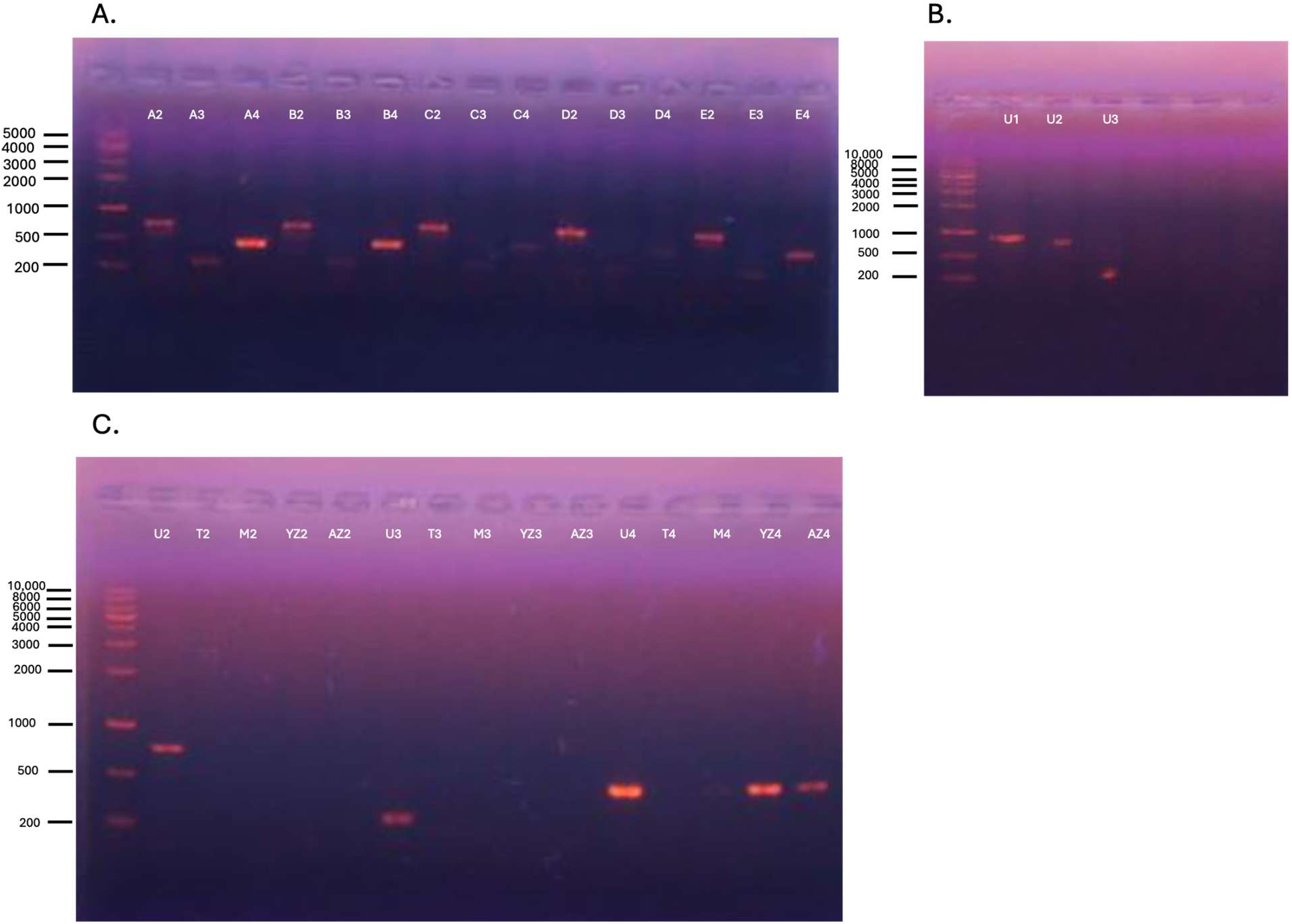
Agarose gel electrophoresis of PCR products from pooled fecal samples. (A) Primer sets; crAss-like02, crAss-like03 and crAss-like04 (expected sizes 752bp, 249bp and 472bp, Table 1). Lanes: 1, 5 kbp ladder; 2-4, composite A; 5-7, composite B; 8-10, composite C; 11-13, composite D; 14-16, composite E. (B) Primer sets: crAss-like01(850bp), crAss-like02 and crAss-like03. Lanes: 1, 10 kbp ladder; 2-4, composite U. (C) Lanes: 1, 10 kbp ladder; 2-6, composites U, T, M, YZ, AZ with primer set crAss-like02 (752bp); 7-11, same composites with set crAss-like03 (249bp); 12-16, same composites with primer set crAss-like04 (472bp). Composite identities are defined in Supplementary Table S13.

**Table 1:**
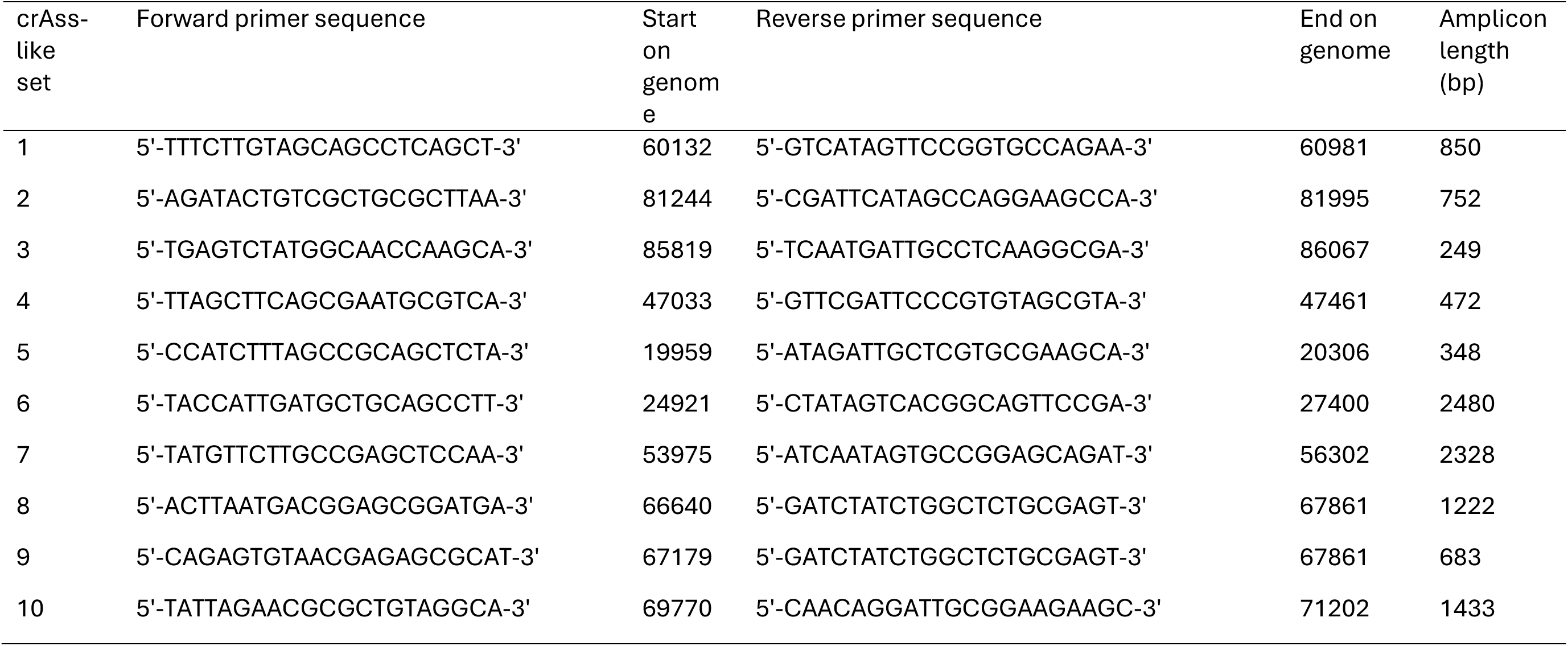
Oligonucleotide primer pairs designed and used in this study.

### 3.6 Taxonomic Analysis Reveals a New Species and Genus

Genome-wide proteomic tree reconstruction clustered the Egyptian phage within a monophyletic group alongside four members of the family *Darmviridae* inside the order *Crassvirales* (Supplementary Figures 6,7). Single-protein Maximum Likelihood phylogenies based on core marker proteins—specifically the major capsid protein (MCP), large terminase subunit (*terL*), portal protein, and DNA primase—along with a concatenated core-proteome tree firmly established its positioning inside the *Darmviridae* family (Figure 5, Supplementary Figure 8)^4^. This placement is further supported by the presence of two signature marker genes, *NC_04SS77 Gp42* and *NC_04SS77 Gp80*, which are conserved in over 99% of all known *Crassvirales* genomes^4^. Moreover, the classification of the Egyptian phage within the new family of *Darmviridae* is also suggested by the shared orthologous percentage (Supplementary Figure 9).

**Figure 5:**
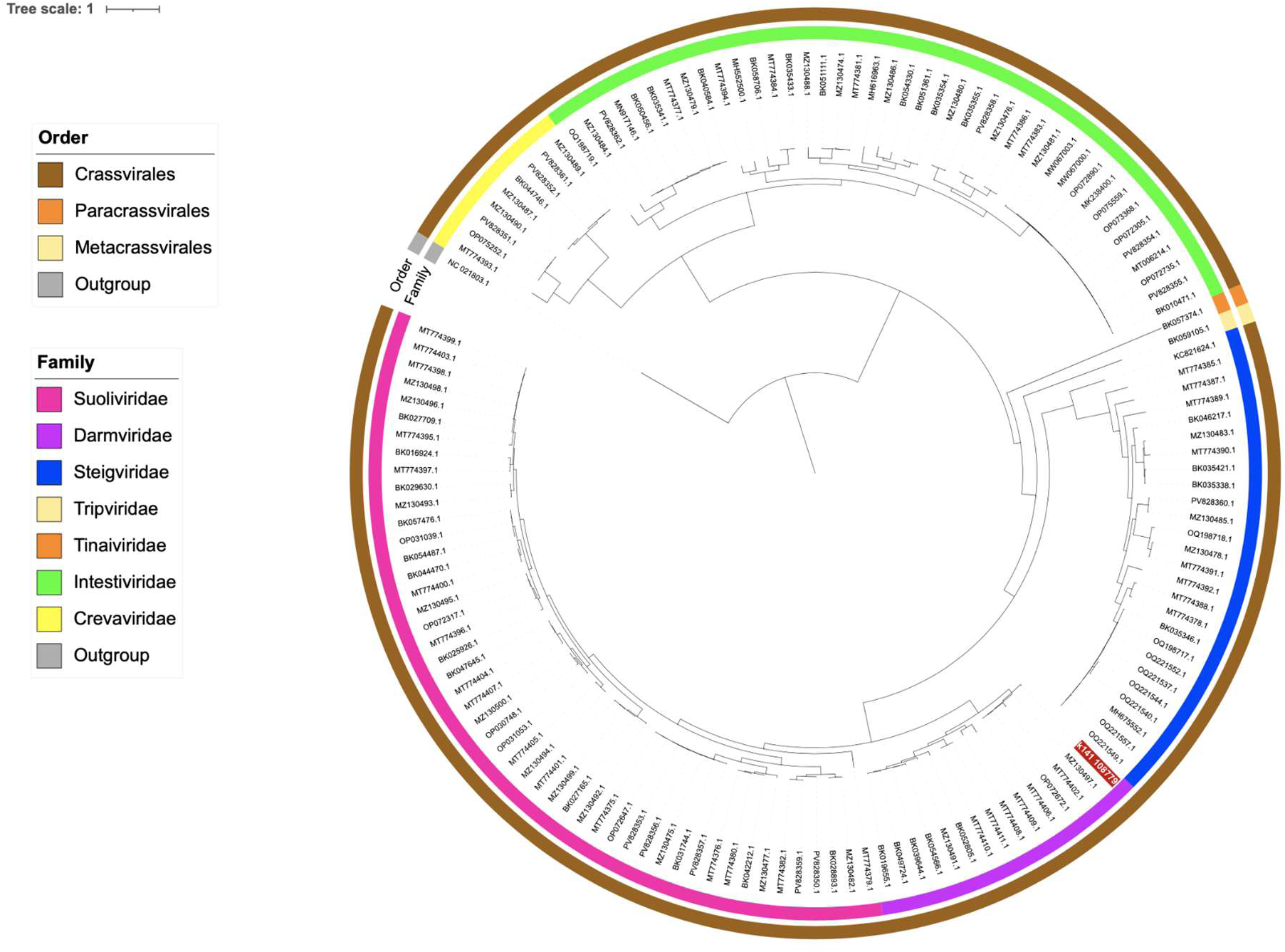
Core-gene phylogenetic tree of the three orders based on concatenated proteins. The concatenation of four gene alignments (major capsid protein, large terminase subunit, portal, and primase) was used to infer a maximum likelihood tree. The Egyptian crAss-like phage is highlighted in red. The outer ring represents order, and the inner ring represents family. The tree is midpoint rooted, with Cellulophaga phage phi13:2 (NC_021803) as an outgroup.

To evaluate the exact taxonomic level of this isolate under ICTV criteria, total average nucleotide identities (tANI) were calculated using VIRIDIC^4,54^. The Egyptian phage shared less than 50% tANI with all currently recognized reference members of the *Darmviridae* family, falling well below the 70% intergenomic threshold established for genus demarcation and indicating that it represents a novel genus (Figure 6).

**Figure 6:** Genomic comparison heatmap of the Egyptian crAss-like phage with members of the family *Darmviridae* and the top six NCBI MAGs hits (starred). The heatmap, generated by VIRIDIC and clustered using the complete linkage method, shows intergenomic similarity values (right half, colour scale from yellow (low) to blue (high)) and alignment indicators (left half). No pair exceeds the 95% total average nucleotide identity (tANI) species demarcation threshold, confirming the Egyptian phage as a new species.

Furthermore, to investigate any distant relationships, we performed BLASTn homology searches of the assembled Egyptian genome against the NCBI nt database. Surprisingly, we found six metagenome-assembled genomes (MAGs) from China, Japan, Madagascar, and Cameroon, covering more than 80% of our phage with varying identities (i.e., 87% to 96%)(Supplementary Figure 10). Despite their high similarities, individual MAG alignments showed that none had an identical nucleotide sequence nor amino acid sequence with the Egyptian crAss-like phage, confirming the fact they all had different genomes (Supplementary Figure 11).

Average amino acid identity (AAI) calculations by EzAAI showed moderate protein similarity of the Egyptian phage to the closest four members of the family Darmviridae (37-69% genome coverage with 82-86% AAI), while higher amino acid identity (59-85% genome coverage with 89-95% AAI) to the top six MAGs (Supplementary Table S15)^43^.

Accordingly, given this high sequence similarity to entries in the NCBI nt database, tANI was also computed between the Egyptian phage and these MAGs (Figure 6). Surprisingly, the Egyptian phage shares more than 70% tANI with all MAGs genomes, but none was similar enough to share >95% tANI, confirming earlier predictions of new species (Figure 6). Consequently, current members of the *Darmviridae* family could be divided into eight putative genera and 15 distinct species, with the Egyptian crAss-like phage likely constituting a novel genus and species.

## 4. Discussion

The primary objective of this study was to investigate the prevalence and genomic diversity of crAss-like phages within an underexplored region, successfully assembling and validating the first complete crAss-like phage genome from an Egyptian population. Phylogenomic characterization of the 101,034 bp circular genome (contig k141_108779) identified taxonomic features aligning it with the family *Darmviridae*, order Crassvirales^4^. Although recent updates by the International Committee on Taxonomy of Viruses (ICTV) established *Darmviridae* as an independent family, it was stated that it can be classified as a subfamily under the closely related and highly compact *Suoliviridae*^4,5^. This may result in similar biological features and evolutionary patterns. The discovery of this phage is significant for multiple reasons: it provides the first insight into the crAss-like phage composition of North African populations, it reveals novel genomic diversity within the *Darmviridae* family, and it highlights a potentially unique non-Westernized viral lineage adapted to the Egyptian gut microbiome.

In silico host prediction models strongly assigned the recently reclassified *Segatella copri* (formerly *Prevotella copri*) as the primary bacterial host, an association validated by multiple protein and sequence-based features including high-confidence sequence matching against the host’s adaptive CRISPR spacer immune pattern^28,67,68^. This finding represents a first-of-a-kind ecological divergence from other crAss-like phages, which predominantly infect members of the genus *Bacteroides*^69^. The dominance of a *Segatella*-infecting phage correlates precisely with the comparative gut microbiome data used in this study, where Shankar *et al.*, highlighted a contrast between the *Prevotella/Segatella*-dominated enterotypes of the Egyptian teenagers and the *Bacteroides*-rich profiles of the matched US candidates, typical for industrialized Western populations^70^. Members of the *Segatella copri* are known to preferentially colonize non-Westernized populations consuming high-fiber, Mediterranean-inspired diets as in Egypt^71–73^. Consequently, the discovery of this novel lineage underscores that the current ICTV reference database for crAss-like phages remains heavily biased toward Eurocentric and North American sampling cohorts.

In accordance with other Crassvirales, the strictly virulent lifestyle predicted for this phage carries important implications for host population dynamics within the human gut ecosystem^74–77^. Functional evaluation of the viral proteome using Random Forest classification identified specific conserved protein domains ^26^. The detection of the cd01123 domain, which belongs to the Rad51/RecA recombinase family, points to an ancient recombination framework capable of interacting with host DNA damage responses^78–80^. In classical temperate models, activated host RecA triggers the self-cleavage of viral repressors to initiate the lytic cycle; however, in a strictly virulent phage, such domains may be co-opted to manipulate host SOS signaling or facilitate autonomous viral recombination during replication^81^. Additionally, the presence of the cd03408 domain, which shares homology with prokaryotic SPFH proteins such as HflK and HflC, indicates a sophisticated interaction with membrane-associated complexes. In *Escherichia coli*, HflK and HflC regulate lysogenic choices by modulating the proteolysis of the lambda CII repressor^82^.In the context of *Crassvirales* biology, which is renowned for maintaining high-abundance, long-term persistence in the human gut without stable lysogenic integration, this domain likely does not function as a standard lysis coordinator^69,74^. Instead, it may serve as a molecular anchor that senses host envelope stress, allowing the phage to shift between an aggressive lytic cycle and a chronic, non-lethal “carrier state” or pseudolysogenic latency^83,84^. This mechanism would enable long-term viral persistence and stable coexistence with the host population without driving it to extinction.

A central finding of this research is genetic codon reassignment. To understand this architecture, it is critical to distinguish between local codon recoding—which represents a context-dependent, mRNA-mediated alteration that directly competes with standard translation, such as programmed ribosomal frameshifting or stop-codon readthrough—and true genetic codon reassignment, which reflects a global, genome-wide, unconditional change in codon meaning^85,86^. The Egyptian phage uses an alternative genetic code, particularly Genetic Code 15, where the canonical TAG *amber* stop codon is reassigned to encode the glutamine (Q) amino acid. This results in an optimization of the predicted genes, increasing coding density to over 90% and restoring full-length functional open reading frames for essential proteins such as Terminase large subunit (TerL), portal proteins and RNA polymerase subunits. However, evidence of functional protein translations must be investigated to confirm phage genetic reassignments^59,87^. Although *Crassvirales* represents one of 8 phage clades known to use stop codon reassignments in human and animal gut microbiomes, this feature appears to occur within specific families, particularly *Steigviridae* (formerly Beta) and *Suoliviridae* (formerly Delta)^5,60,88^.

The genomic organization of the Egyptian phage is consistent with the bi-*amber* domains proposed by Ivanova *et al.*^65^ and further described by Borges *et al.* and confirmed via metaproteomics by Peters *et al.* ^59,60^. This bi-modal distribution of TAG amber stop codons suggests a sophisticated temporal regulation of the viral life cycle. The low-*amber* domain constitutes early-stage genes involved in metabolism, DNA replication, and transcription. This domain avoids in-frame reassigned TAG codons, allowing genes to be translated immediately with the standard code of the host’s machinery. Curiously, these early genes produce nearly identical proteins when translated by either Genetic Code 11 or 15. This concept of “dual-coded” blocks was hypothesized by Pfennig *et al.* as a ‘soft switch’ strategy in which phages continue the translation of essential early genes uninterrupted by changes in the translation environment^89^. Meanwhile, the high-amber domain contains late-stage infection genes such as tail fiber and tail adaptor proteins, phage stabilization protein, terminase large subunit, and portal protein in addition to Mannosyl-glycoprotein endo-beta-N-acetylglucosaminidase enzyme and multiple proteins of unknown function. These proteins would fit into three of the four gene families of crAss-like phages described by Borges *et al.* to preferentially use TAG over CAG and CAA to encode glutamine. These families are the tail tube gene family, the lysozyme amidases, and the gene family of unknown function^60^. Thus, late genes would be biologically meaningful if the translation apparatus recognizes *amber* TAG as a codon for glutamine (Q). It is now believed that this timed expression of late assembly and packaging proteins regulates host lysis and prevents premature expression of key structural genes.

Three cases of *amber*-reassigned phage sequences have been reported in a large-scale metagenomic study of stop codon reassignments, which identified five CRISPR spacers identical to three TAG-reassigned phage genomes. Interestingly, these bacteriophages infect two standard-coded *Prevotella* strains isolated from different human sites^65^. Another *amber* reassignment has been documented in all lineages of the widespread lytic Lak megaphages to date^40,90^. These megaphages infect *Prevotella* species as well, including Segatella copri (formerly Prevotella copri)^87^. Moreover, an assembled amber-reassigned prophage was integrated into a standard-coded *Prevotella* contig^60^. The observation of stop codon reassignments in these phages raises questions about the relationship with their target hosts. A large-scale evolutionary genomic survey on more than 9,000 phage genomes observed the prevalence of reassigned phages with members of the standard-coded Firmicutes and Bacteroidetes^60^. This suggests that alternative genetic code may be more evolutionarily dynamic and host-associated than previously recognized.

Surprisingly, by exploring the reassignments among the functional categories of the genes, we confirmed that the gene encoding major capsid protein (MCP) shows no alternative codon usage. Capsid proteins, although they show low sequence conservation, are the most prevalent of *Crassvirales* genes, making them the most suitable markers for reconstructing phylogenetic trees ^91^. This could be explained by diversifying selection pressure, increasing trait differences and potentially leading to speciation. Meanwhile, since capsid genes are the most affected by host translational bias ^92^, they are under strong pressure to adapt their codon usage to their host’s translation preferences. This points to the importance of their efficient translation in phage vitality.

Alterations of the translation mode within the phage life cycle are mediated by a “code change module” located upstream of the lysis cassette and composed of a suppressor tRNA, a corresponding aminoacyl-tRNA synthetase (aaRSs) and/or a release factor (RF1 or RF2) ^60,89^. These components are used during transcription of alternatively coded genes within subsequent structural and lysis genes ^60^. The Egyptian phage encodes four tRNAs, none of them with the anticodon CUA required to suppress the standard stop codon TAG. However, absence of suppressor tRNAs was recorded in a reassigned phage of the *Suoliviridae* family ^5^ and has been reported for other reassigned phages ^65^. The presence of codon reassignments without a detectable suppressor tRNA suggests that these phages either rely on host-encoded suppressors, encode non-canonical computationally-unrecognized suppressors, or have incomplete phage genome assembly. However, annotation of the Egyptian genome reveals the presence of a glutamyl-tRNA(Gln) amidotransferase subunit, suggesting an alternative enzymatic route capable of supporting indirect tRNAGln charging in the absence of glutaminyl-tRNA synthetase ^93^. Similar to other *Crassvirales,* the Egyptian crAss-like genome lacks any encoded release factors.

Taxogenomic and comparative genomic analyses demonstrate that this isolate represents a distinct evolutionary branch within the *Darmviridae* family. Although proteome-based trees generated via ViPTree and single-protein Maximum Likelihood phylogenies group the Egyptian phage alongside established genera such as *Smehivirus*, *Sdahravirus*, and *Burzaovirus*, its total intergenomic similarity falls well below standard taxonomic boundaries. Pairwise total average nucleotide identities (tANI) calculated via VIRIDIC revealed that the phage shares less than 50% identity with all recognized type species, significantly below the 70% threshold designated by the ICTV for genus demarcation. Furthermore, while homology searches against the NCBI *nt* database identified six closely related metagenome-assembled genomes (MAGs) from Japan, China, Madagascar, and Cameroon sharing up to 96% identity across limited regions, pairwise tANI comparisons with these global sequences did not exceed 87%. Because none of these relatives surpass the 95% tANI boundary required for shared species status, these data confirm that the Egyptian phage represents both a novel species and a novel genus, supporting prior hypotheses regarding the high degree of genomic heterogeneity hidden within individual *Crassvirales* families^5,77^.

Population surveillance across 252 individual Egyptian fecal samples successfully confirmed the real-world circulation of this computationally discovered viral lineage. The variable detection frequencies observed between the designed primer sets—where the crAss-like04 locus was widely prevalent across eight composite pools while crAss-like01 was captured in only one— highlight potential issues with primer efficiency or the existence of micro-heterogeneous strains circulating locally. These sequence variations among amplicons, such as the U1 amplicon sharing only 75.7% identity with reference repositories, likely reflect active selective pressures, including host defense mechanisms and dietary fluctuations unique to the North African gut virome^21,94,95^.

Despite the robust combination of metagenomic assembly, phylogenomic profiling, and molecular validation, several limitations must be acknowledged. Deriving the viral genome entirely from metagenomic read sets rather than a cultured isolate leaves open the possibility of minor assembly gaps or the omission of non-canonical elements. Additionally, the pooling of fecal samples into composite groups masks individual-level variations in viral abundance, and a high proportion of the predicted genes remain categorized as hypothetical proteins with uncharacterized functions. Nevertheless, this study provides a complete genomic foundation for a novel *Crassvirales* genus. The identification of a strictly lytic lifecycle, combined with a unique modular architecture adapted to a *Segatella copri* host, positions this phage as a strong candidate for population-specific microbiome biomarkers and targeted phage-based therapeutic applications designed to modulate the human gut virome. Future long-read sequencing (e.g., Nanopore/PacBio) may help resolve assembly gaps, and metaproteomics could confirm the translation of reassigned genes.

